# Genomic characterization of sub-populations in human pluripotent stem cell-derived retinal progenitor cells that drive retinal layer structure

**DOI:** 10.1101/2025.06.20.660659

**Authors:** Yasuaki Iwama, Tomohiro Masuda, Mika Yoshimura, Mikiya Watanabe, Martin Friedlander, Kohji Nishida, Masayo Takahashi, Itoshi Nikaido, Mchiko Mandai

## Abstract

The mechanism underlying retinal spheroid layer formation was investigated using Rax::GFP-positive retinal progenitor cells from human embryonic stem cell-derived retinal organoids. Single-cell RNA sequencing revealed that well-layered spheroids transiently activated canonical WNT2B–FZD7 signaling followed by temporary expression of non-canonical WNT5A. Despite structural differences in vitro, non-layered and well-layered retinal spheroids on differentiation Day 60 successfully developed into cone and rod subtypes of photoreceptors and established synaptic connections with host bipolar cells after transplantation in a retinal degeneration rat model, resulting in light-evoked functional recovery.

Additionally, part of the presumable Rax::GFP-positive retinal progenitor cells differentiated into non-retinal lineages, including ciliary marginal zone-like, retinal pigment epithelium, and spinal cord-like tissues in vitro, reflecting the developmental plasticity of RAX-positive cells. These findings suggest that canonical and non-canonical WNT signaling pathways sequentially orchestrate early retinal morphogenesis, whereas environmental factors within the host retina strongly drive the alignment and functional integration of graft photoreceptors.

## INTRODUCTION

Cell-based regenerative therapy is considered a promising strategy for restoring visual function in retinal degenerative diseases, such as retinitis pigmentosa. ^1,2,3^ In a recent first-in-human trial for patients with retinitis pigmentosa, our team used retinal sheets excised from human induced pluripotent stem cell-derived retinal organoids (ROs) and showed stable graft survival and safety two years after transplantation; ^4^ however, improvements in visual function remain ambiguous. For high visual function, such as fine-tuning of contrast and motion, precise organization of retinal cells, and well-aligned synapse connectivities between the grafted photoreceptors and the inner retina of the host are required. ^5^ Among the various signaling pathways associated with retinal layer structuring during mammalian retinal development, including NOTCH, WNT, HH, and FGF, the canonical WNT pathway is a key candidate for early retinal development. ^6,7,8,9^

In previous studies, mouse retinal progenitor cells (RPCs) in the early stage, equivalent to differentiation Day (DD) 25–30 in human RPCs, have been reported to develop inner and outer nuclear layers more frequently after allotransplantation than those in later stages. ^10,11^ This suggests that ROs at this stage include cells that control retinal layer formation. Among young ROs, RAX, an RPC marker, is an essential transcription factor that guides retinal development. ^12,13^ Intriguingly, when we sorted the RAX-positive and SSEA1-negative (removing ciliary margin cells or immature mitotically active cells) populations from the dissociated RO cells on DD25–30, ^14,15,16^ two different protocols of RO induction by bone morphogenetic protein 4 (BMP4) treatment on either DD3 or DD6 resulted in three distinct characteristi phenotypes in re-aggregated spheroids from the sorted cells: non-layered circular retinal spheroids (NL- spheroids), well-layered circular retinal spheroids (WL-spheroids), and polarized spheroids (P-spheroids).^17^ Although RAX-positive cells in ROs on DD25–30 are generally regarded as RPCs, ^12,13^ our preliminar data suggest that these cell populations exhibit notable heterogeneity, including sub-populations likely involved in retinal laminar structures.

The aim of this study was to better characterize cellular heterogeneity within RPCs in ROs at an early stage of differentiation and to identify sub-populations that drive retinal layer structures. We investigated the gene expression patterns of the composite cell populations of these three types of spheroids derived from the Rax::Venus human embryonic stem cell (ESC) line using single-cell RNA-seq (scRNA-seq) analysis. We identified possible sub-populations of Rax::Venus-positive progenitor cells that 1) contribute to the formation of retinal layer structures and 2) differentiate into non-retinal cells. Since structured RO integration seems to promote stable graft survival as well as post-transplantation function, we also evaluated whether RPC spheroids with or without layered structures may lead to different outcomes after transplantation in rats with end-stage retinal degeneration.

## RESULTS

### Characteristic spheroid morphologies are obtained based on Venus-intensity and SSEA1-absence

To detect the variety of presumable RPC sub-populations, we first investigated the spheroid phenotype by re-aggregating the Rax::Venus-positive cells from Rax::Venus ESC line (KhES-1)-derived ROs on DD25 using Y-27632 (ROCK inhibitor), 3 μM CHIR99021 (GSK3β inhibitor), SAG (smoothened agonist), and FGF8 (fibroblast growth factor). Since RAX expression is considered an appropriate RPC marker at this stage, ^12,17^ we defined RPCs in ROs as Rax::Venus-positive and SSEA1-negative cells to exclude ciliary margin cells or immature mitotically active cells. ^14,15,16^ ROs were differentiated via a modified serum-free floating culture of embryoid body-like aggregates using the quick aggregation (SFEBq) method and BMP4 treatment. ^15,18^ During differentiation, two different timings of BMP4 treatment at DD3 or DD6 yielded ROs with different profiles of Venus-intensities in Rax:: Venus-positive cells (D3-BMP4 and D6-BMP4, respectively), whereas the absence of BMP4 treatment did not produce Rax::Venus- positive ROs (no-BMP4)(Figures 1*A–1C)*. Then, the D3- and D6-BMP4 ROs on DD25 were dissociated into single cells and sorted based on Venus-intensities with another parameter of SSEA1-absence using fluorescence-activated cell sorting (FACS).

**Figure 1.**
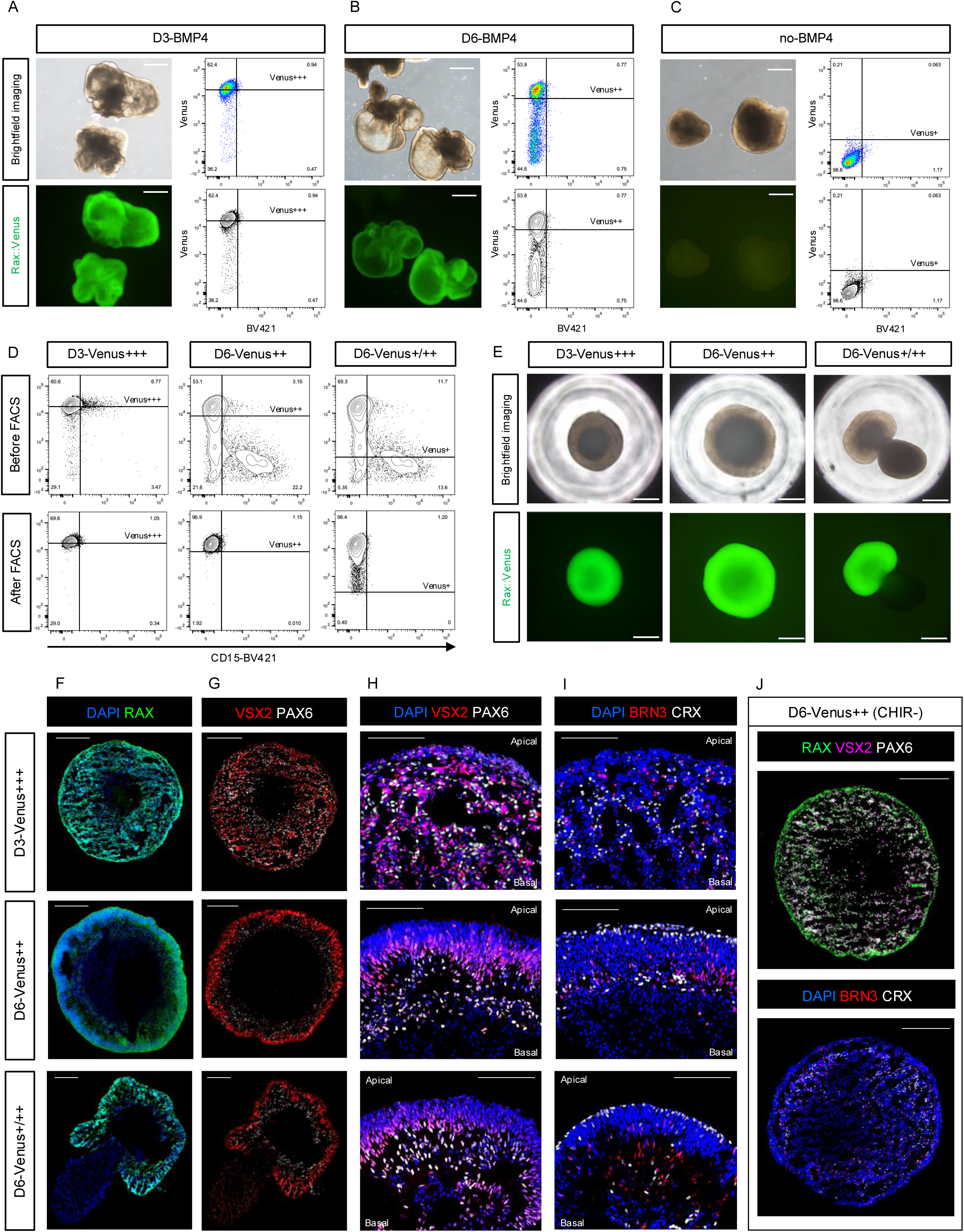
Characteristic spheroid morphologies after fluorescence-activated cell sorting (FACS)- sorted progenitor cell re-aggregation. (A–C) Phenotypes and flow cytometry profiling of D3-BMP4 (A), D6-BMP4 (B), and no-BMP4 organoids (C). D3- and D6-BMP4 organoids were those subjected to BMP treatment on DD3 and DD6, respectively. No-BMP4 organoids were not subjected to BMP treatment. Top right: Venus-A and BV421-A profiles in; bottom right: BV421-A and Venus-A profiles in contour plots. We set the Venus+++, Venus++, and Venus + lines by referrng to the contour plot of Venus-intensity profiles of D3-, D6-, and no-BMP4 organoids, respectively (A–C). (D) Venus-A and CD15-BV421-A profiles of D3-Venus+++ sorted cells, D6-Venus++ sorted cells, and D6- Venus+/++ sorted cells. FACS was conducted based on Venus-intensities, gating for BV421-negative cells. Top: profiles before FACS; bottom: profiles after FACS. (E) Characteristic spheroid morphologies after re-aggregation of D3-Venus +++, D6-Venus++, and D6- Venus+/++ FACS-sorted cells. (F–I) Immunostaining images of each spheroid on DD60–70 with Rax::Venus (Green, F) and antibodies against VSX2 (Red, G and H), PAX6 (White, G and H), BRN3 (Red, I), CRX (White, I), and DAPI (Blue, F, H and I). (J) Immunostaining images of spheroids derived from D6-Venus++ cells in DD60-70 without CHIR99021 treatment. VSX2 (magenta, top), PAX6 (white, top), BRN3 (red, bottom), and CRX (white, bottom) were expressed in scattered cells. Scale bars: 500 μm (A–C, E), 200 μm (F, G, J), and 100 μm (H, I).

Interestingly, D3-Venus+++, D6-Venus++, and D6-Venus+/++ formed different types of spheroids based on their morphology and Venus expression patterns (Figures 1*D* and 1*E*). On DD60, 35 days after re- aggregation, D3-Venus+++ cells developed characteristic spheroids with relatively small Rax::Venus-positive non-layered structures (NL-spheroid, hereafter, left panel in Figure 1*E*), whereas D6-Venus++ cells developed into relatively large spheroids consisting of Rax::Venus-positive well-aligned layer structures (WL-spheroid, hereafter, middle panel, Figure 1E). Although we sorted the RAX-positive cells, the D6-Venus+/++ cells developed into polarized spheroids consisting of Rax::Venus-positive and - negative parts (P-spheroid, hereafter, right panel, Figure 1*E*). Immunohistological examination showed that NL-spheroids contained cells positive for VSX2, PAX6, CRX, and BRN3, suggesting that they were differentiating into retinal tissue but did not form retinal layer structures (tops, Figures 1*F*–1*I*). In contrast, WL-spheroids and the RAX-positive portion of the P-spheroids revealed neural epithelium-like structures aligned with VSX2-positive cells apically and PAX6-positive cells basally, with the presence of CRX-positive photoreceptor precursor cells predominantly apically and BRN3-positive retinal ganglion cells basally (middles and bottoms, Figures 1*F*–1*I*). Additionally, D6-Venus++ cells developed into NL- spheroids without CHIR99021 treatment in four independent trials (Figure 1*J*). These results indicate that 1) NL-spheroids lack cell populations controlling retinal layer structures, 2) D6-Rax::Venus+/++ contains cells other than RPCs, and 3) the canonical WNT pathway is involved in the formation of retinal layer structures in WL-spheroids.

### Overall cell populations were similar between NL- and WL-spheroids, while P-spheroids contained three characteristic cell populations

To evaluate the cell populations comprising these three different spheroid patterns, we conducted a scRNA-seq analysis of the merged data of these spheroid types at four time points during differentiation (Figure 2*A*). First, we focused on the final phenotype of each spheroid pattern to investigate differences in composite cell populations. On post-sorting Day 35 (PS-D35, DD60), no spheroid pattern-specific cell populations were observed between NL- and WL-spheroids (Figure 2*B*). At this time point, RPCs (RAX +, VSX2 +), mitotic cells (S and G2/M phases), transitional precursor cells (ATOH7 +), retinal ganglion cells (RGC, SNCG+ and POU4F2 +), amacrine cells (AC, TFAP2A +), horizontal cells (HC, ONECUT1 +), and photoreceptors (PR, CRX + and RCVRN+) were similarly observed in NL- and WL-spheroids ^19,20^ (Figures 2*C and* 2*D*).

**Figure 2.**
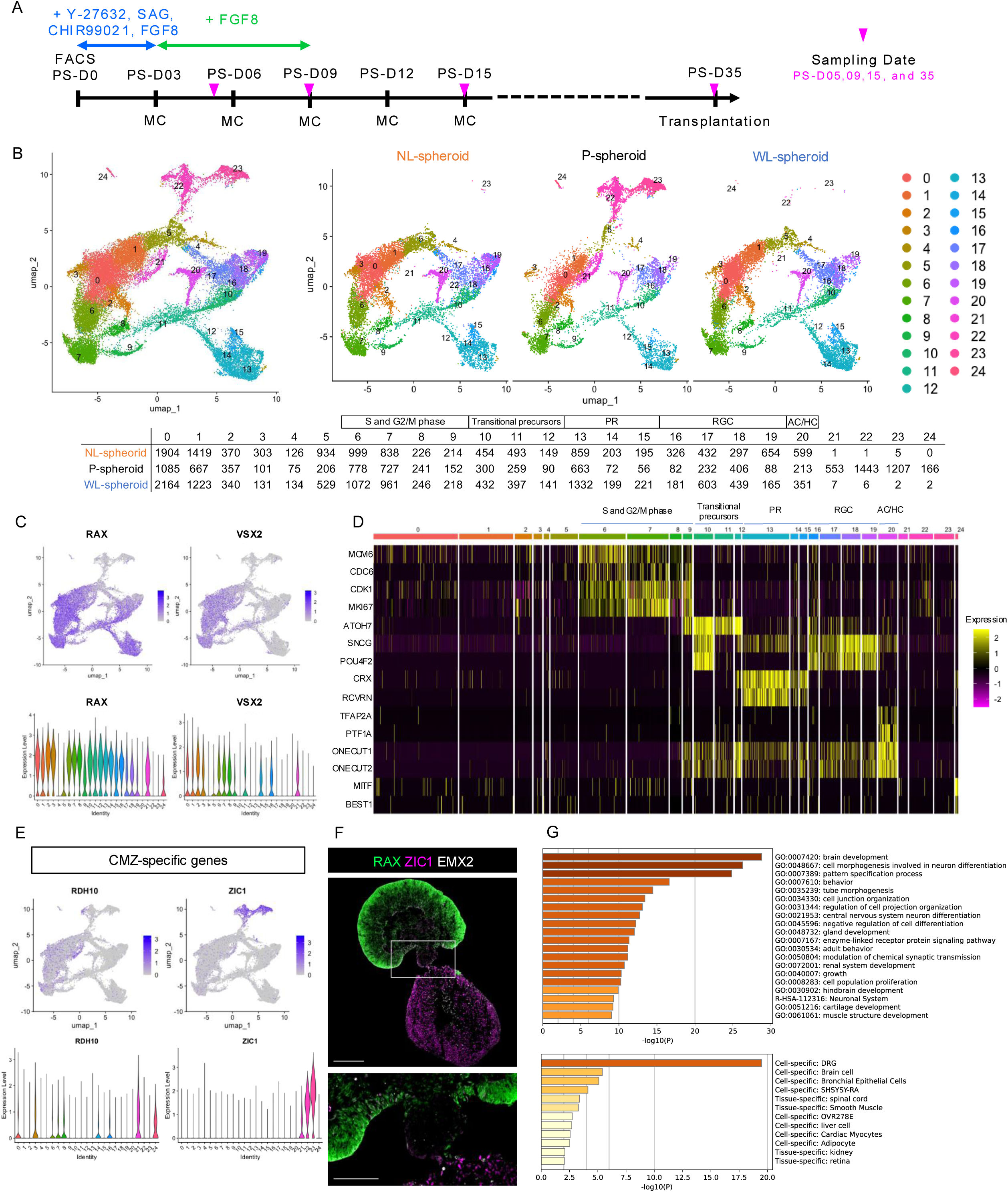
Single-cell RNA-seq analysis of three characteristic spheroids on PS-D35. (A) Differentiation protocol highlighting the required reagents and the sampling time points. (B) UMAPs of NL- (D3-Venus+++), WL- (D6-Venus++), and P- (D6-Venus+/++) spheroids on PS-D35. (C) Feature plots showing the expression of eye field transcriptional factors (RAX and VSX2). (D) Markers characteristic of retinal ganglion cells (SNCG and POU4F2), photoreceptors (CRX and RCVRN), amacrine/horizontal cells (TFAP2A and ONECUT1), and retinal pigmental epithelium populations (MITF and BEST1). (E) Feature plots (top) and violin plots (bottom) showing the expression of genes associated with the ciliary marginal zone. Cluster 21, which is specific in P-spheroid, includes cells weakly positive for RDH10 and ZIC1. (F) Immunostaining images of P-spheroid on DD60–70. Rax::Venus (green)- and ZIC1 (magenta)-positive cells were observed at the peripheral retina. (G) Gene ontology analysis of 500 top-ranked marker genes of P-spheroid-specific RAX-negative clusters (clusters 22 and 23) using Metascape. Bar graph of enriched terms across input gene lists (top) and summary of enrichment analysis in PaGenBase (bottom). Scale bars: 200 μm (E, top) and 100μm (E, bottom). SAG, smoothened agonist; FGF, fibroblast growth factor; NL-spheroid, non-layered spheroid; P-spheroid, polarized spheroid; WL-spheroid, well-layered spheroid; PR, photoreceptor cell; RGC, retinal ganglion cell; AC, amacrine cell; HC, horizontal cell.

In contrast, three phenotype-specific cell populations were observed in the P-spheroids: 1) cluster 21, 2) clusters 22 and 23 (RAX-negative population), and 3) cluster 24 retinal pigment epithelium (RPE, MITF +, and BEST1 +) (Figure 2*B* and 2*G*). In the scRNA-seq analysis, cluster 21 expressed RAX, RDH10, and ZIC1, which have previously been reported to be ciliary marginal zone (CMZ) marker genes (Figure 2*E*). ^15^ We considered cluster 21 as a CMZ-containing retinal cell population because RAX- and ZIC1-positive cells were observed in the peripheral retinal area outside the EMX2-positive region in an immunohistological image (Figure 2*F*). Clusters 22 and 23 contained significantly upregulated differentially expressed genes (DEGs) (Wilcoxon rank-sum test, Supplemental File 01). Gene ontology analysis, performed with Metascape using the top 500 upregulated DEGs based on average log2 fold change, demonstrated that this cell population was similar to those associated with brain development and neuronal differentiation, which are specific to dorsal root ganglion cells, brain cells, and spinal cord-like tissues (Figure 2*G*).

### RPC sub-populations with the canonical WNT receptor FZD7 potentially guide the self-organization of retinal layer structures

Second, we focused on the time point PS-D05 before the phenotypic differences between NL- and WL- spheroids became evident, which was PS-D09 (Figures 3*A* and 3*B*). Merged scRNA-seq data of NL- and WL-spheroids on PS-D05 consisted of 16 clusters with almost one group of RPC clusters, which expressed eye field transcriptional factors (Figures 3*C* and 3*D*). ^21^ To infer intercellular communication (ligand-receptor interactions), we used CellChat on these RPC clusters to detect significant signaling pathways. ^22^ We identified 68 potentially significant pathways, including the WNT, NOTCH, HH, and FGF pathways (Figure S1). As for the WNT pathway, clusters 3 and 13 expressed ligands and WNT2B-ligand associated pathways were detected as having a high contribution (Figure 3*E* and 3*F*). Both clusters were WL-spheroids-dominant, but there appeared to be no obvious differences between NL- and WL-spheroids in the violin plot (Figure 3*G*).

**Figure 3.**
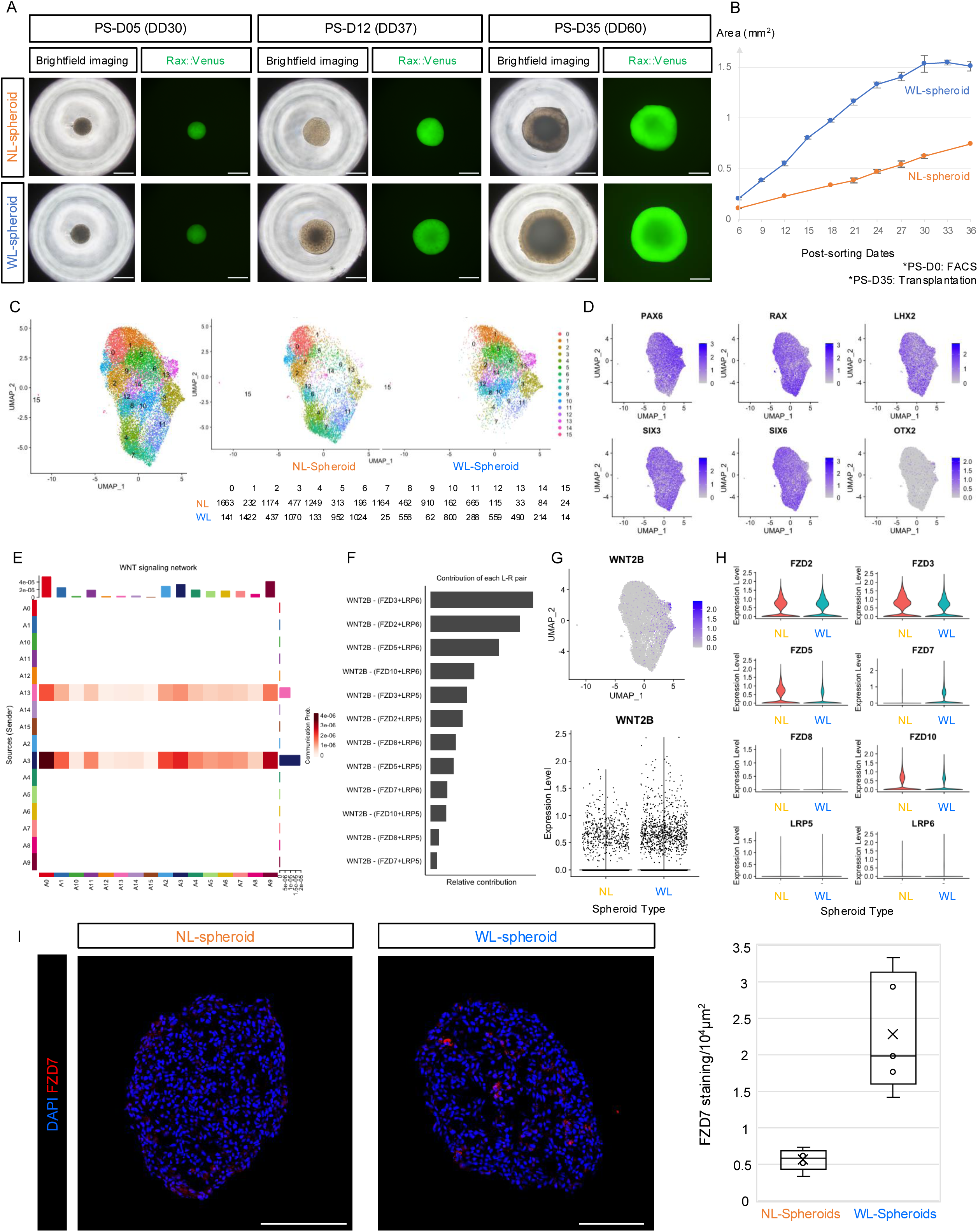
Single-cell RNA-seq analysis of Non-layered (NL-) and Well-layered (WL-) spheroids on PS-D05. (A) Time-lapse imaging of representative cases of NL- and WL-spheroid (Top and bottom, respectively). (B) Temporal changes in the area of each spheroid in brightfield images. Red and blue meant NL- and WL-spheroid, respectively. (n = 4, mean ± SD). (C) UMAPs of NL- (D3-Venus+++) and WL- (D6-Venus++) spheroids on PS-D05. (D) Feature plots showing the expression of eye field transcriptional factors in blue. (E) Heatmap of the cellar communication of each cluster associated with the WNT pathway. (F) Relative contribution of each ligand-receptor pair to the overall communication network of the WNT signaling pathway. Among all known ligand-receptor pairs, WNT2B ligand-associated pathways were detected as ones with high contribution. (G) Feature plot (top) and violin plot (bottom) showing the expression of WNT2B. Cells in clusters 3 and 13, more common in the WL-spheroid, tended to express WNT2B. (H) Violin plots showing the expression of receptor genes associated with the WNT signaling pathway. (I) Immunostaining images of NL- and WL-spheroid on PS-D05. Quantification is presented as box plots. Each dot represents biologically independent, each x-mark the mean value, and each line the median [n = 5 in each group; *p* = 0.0081 (95% confidence interval: −2.74 to −0.73) in Welch’s t-test]. Scale bars: 500 μm (A) and 100 μm (I). NL-spheroid, non-layered spheroid; WL-spheroid, well-layered spheroid.

Considering Figure 1*H* results and the fact that CHIR99021 treatment enhances all canonical WNT pathways, ^23^ we focused on the receptor side. Among all WNT-related receptors, FZD7 was more highly expressed in WL-spheroids than in NL-spheroids, unlike the other canonical WNT receptors, in our scRNA-seq analysis (Figure 3*H*). Immunohistochemical images of PS-D05 demonstrated that significantly more FZD7-positive cells were observed in WL-spheroids than in NL-spheroids (*p* < 0.01, Welch’s t-test, Figure 3*I*). We tested whether human recombinant WNT2B affected the spheroid phenotype of D6- Venus++ cells from PS-D0 to PS-D6 in the absence of CHIR99021; however, these cells still differentiated into NL-spheroids. We still need to optimize these experiments, but the requirement for CHIR99021 collectively suggests that the WNT2B-FZD7 pathway potentially guides the formation of retinal layer structures (Figures 1*H* and 3*F–I*).

### Non-canonical WNT5A is potentially associated with the self-organization of retinal layer structures following WNT2B-associated pathway activation

Third, we focused on the time points of PS-D09 and PS-D15 when phenotypic differences between NL- and WL-spheroids began to form (Figure 3*A*). On PS-D09, the merged scRNA-seq data of NL- and WL- spheroids consisted of two groups of clusters, RPCs and precursors (Figure S2), and there were no differences in the precursor cell populations between these spheroids. Therefore, RPC cell populations were a subset, and downstream analysis was performed again with UMAP embedding of min.dist = 0.01 (Figure 4*A*). In this re-clustering of the RPC populations, the right side of the cell clusters (clusters 0, 3, 4, 6, 7, 9, and 10) expressed S- and G2/M-phase marker genes (Figure 4*B*). At this time point, the canonical WNT pathway was detected in CellChat as a significant pathway (Figure S1), but only WNT3A-associated pathways, and not WNT2B-associated pathways, were identified as significant (Figures 4*C* and 4*D*). In contrast, non-canonical WNT (ncWNT) pathways, which were not significant in PS-D05, were significant in PS-D09 (Figure S1). In CellChat, WNT5A-ligand-associated pathways were detected as those with a high contribution to the overall communication network of the ncWNT pathway, mainly in clusters 3, 8, 9, and 12 (Figures 4*E*, 4*F*, and 4*G*). In these clusters, obvious differences were observed in the violin plot between NL- and WL-spheroids (bottom left, Figure 4*G*). Immunohistochemical images of PS-D09 demonstrated that WNT5A-positive cells were detected on part of the surface of NL-spheroids and throughout the surface of WL-spheroids (Figure 4*H*).

**Figure 4.**
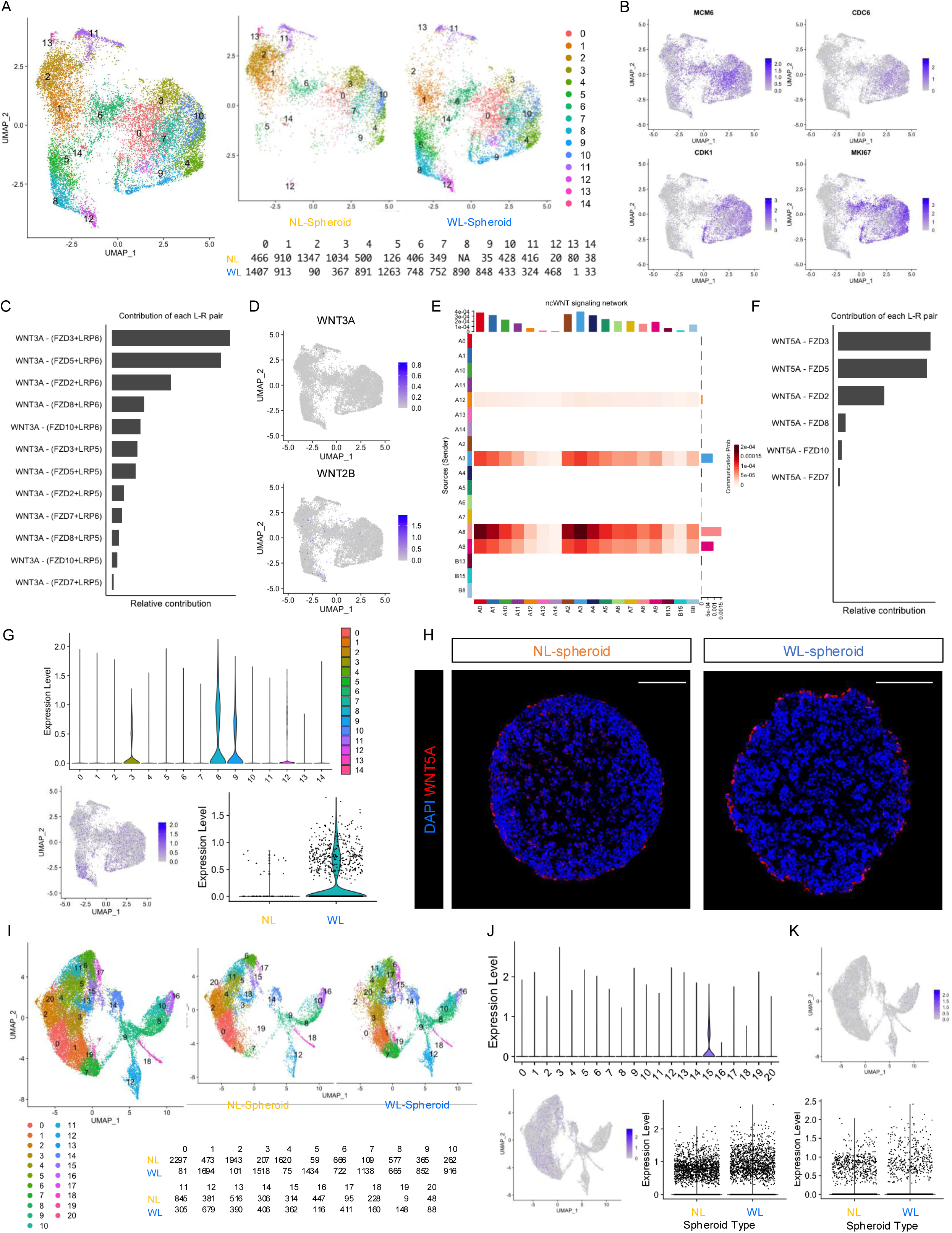
Single-cell RNA-seq analysis of Non-layered (NL-) and Well-layered (WL-) spheroids on PS-D09 and PS-D15. (A) UMAPs for progenitor cell population of NL- (D3-Venus+++) and WL- (D6-Venus++) spheroids on PS- D09. (B) Feature plots on PS-D09 showing the expression of cell division-associated genes (S and G2/M phases) in blue. (C) The expression distribution of signaling genes involved in the inferred significant canonical WNT signaling network on PS-D09 using CellChat. (D) Feature plots showing the expression of WNT3A and WNT2B on PS-D09. (E) Heatmap of the cellar communication of each cluster associated with the non-canonical (ncWNT) pathway on PS-D09. In this analysis, precursor cell clusters (B8, B13, and B15 in Figures S1 and S2) were included. (F) Relative contribution of each ligand-receptor pair to the overall communication network of the WNT signaling pathway on PS-D09. Among all known ligand-receptor pairs, WNT5A ligand-associated pathways were detected as ones with high contribution. (G) Violin and feature plots showing the expression of WNT5A on PS-D09. Cells that express WNT5A were obviously observed in clusters 3, 8, 9, and 12 (top and bottom, left). As for the violin plot of these clusters (bottom, right), WNT5A-positive cells were highly detected in WL-spheroids. (H) Immunostaining images of NL- and WL-spheroids on PS-D09. WNT5A-positive cells were observed on the partial surface of NL-spheroids and on the total surface of WL-spheroids. (I) UMAPs of NL- (D3-Venus+++) and WL- (D6-Venus++) spheroids on PS-D15. (J) Violin and feature plots showing the expression of WNT5A on PS-D15. Cells that express WNT5A were observed in cluster 15 (top and bottom, left). No obvious changes in WNT5A expression were detected between NL- and WL-spheroids (bottom, right). (K) Feature plot (top) and violin plot (bottom) showing the expression of WNT2B on PS-D15. Scale bars: 100 μm (H). NL-spheroid, non-layered spheroid; WL-spheroid, well-layered spheroid.

In contrast, merged scRNA-seq data of NL- and WL-spheroids on PS-D15 demonstrated that WNT5A- positive cells were observed in cluster 15, one of the RPC populations; however, this cluster was not dominant in either NL- or WL-spheroids (Figures 4*I* and 4*J*). No obvious differences in WNT5A expression were observed between phenotypes in the violin plot (Figure 4*J*). Additionally, no clusters clearly expressed WNT2B in the PS-D15. These results suggest that there are RPC sub-populations that temporarily secrete WNT5A after WNT2B activation around PS-D09 in WL-spheroids. However, the addition of the human recombinant protein WNT5A to D3-Venus+++ cell-derived NL-spheroids from PS- D06 to PS-D12 did not induce layer formation (three independent trials).

### Retinal spheroids with or without retinal layer structures showed similar structural and functional engraftment in retinal degeneration rats

Finally, we transplanted NL- and WL-spheroids on DD60, as in previous studies, ^3,10,17^ to 6-month-old SD- Foxn1 Tg(S334ter)3LavRrrc nude rats, which are immunodeficient end-stage retinal degeneration model rats. ^24^ NL- and WL-spheroids were cut into retinal sheets and transplanted subretinally in these rats; to identify the graft-driven light-evoked responses in transplanted retinas, multi-electrode array (MEA) recordings were performed using isolated retinas 300–330 days after transplantation (Figure 5*A*). Non-transplanted eyes were used as controls and retinal ganglion cell (RGC)-derived light-evoked spike counts, the RGC light-evoked responses, were evaluated as described previously. ^3^ Restoration of light responsiveness in host ganglion cells was observed in the grafted area in two of the three eyes transplanted with each spheroid type-derived retinal sheet (Figures 5*B–*5*D*).

**Figure 5.**
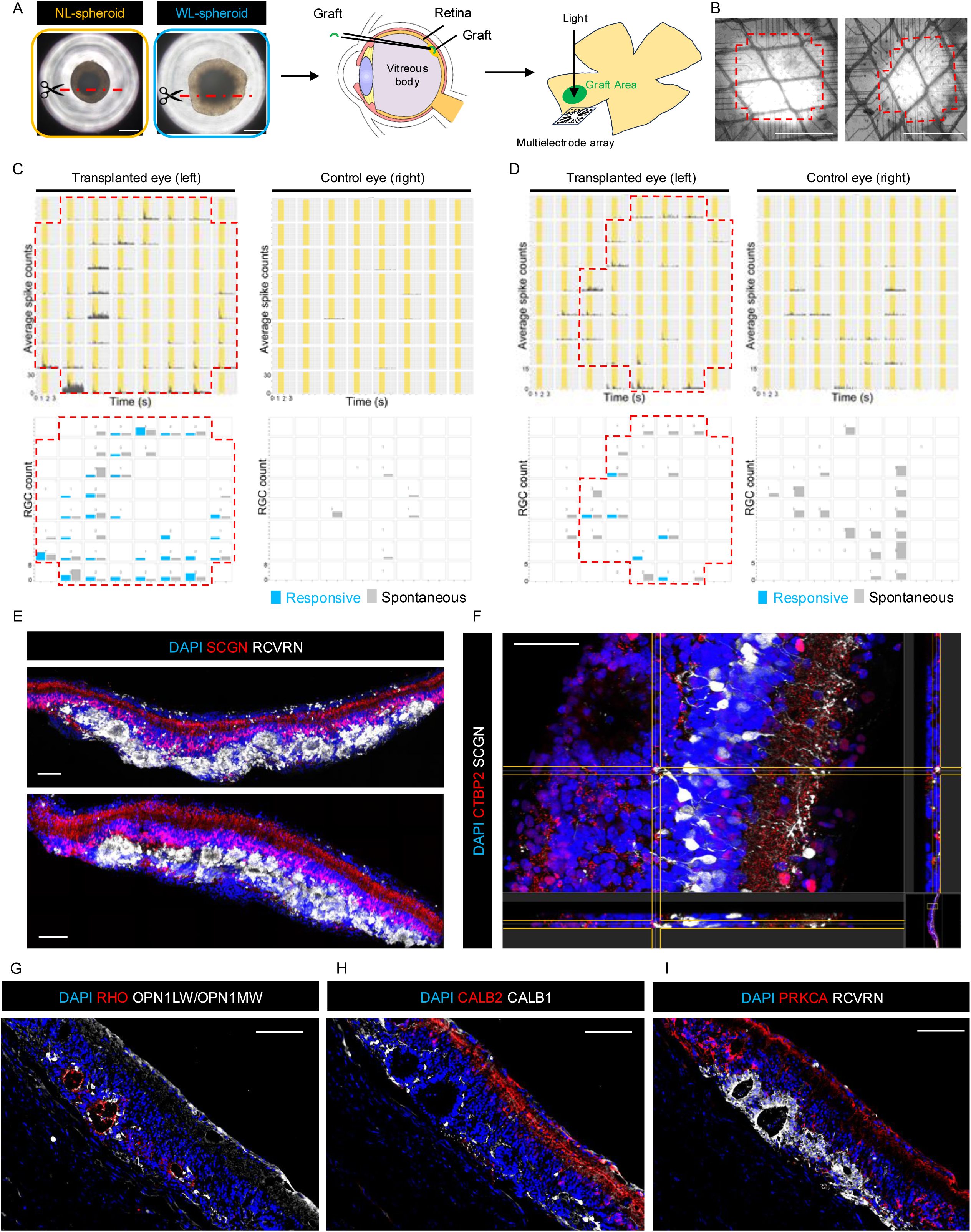
Electrophysiological and histological examination of the graft after transplantation of NL- and WL-spheroids. (A) Schematic illustration summarizing the transplantation in the present study. NL- and WL-spheroids were cut and transplanted subretinally to the retinal degeneration model nude rats on DD60. Electrophysiological examinations using the multi-electrode array (MEA) were performed 300–330 days after transplantation. (B) Representative images of two transplanted rat retinas mounted on MEA electrodes during recording. The electrode positions correspond to the graft area, and the left and right images correspond to Figures 5C and 5D, respectively. (C and D) Results of multi-electrode array (MEA) analysis of the rat retina after transplantation of retinal graft from NL- (C) and WL-spheroid (D). Left and right panels correspond to the transplanted (left eye) and non-transplanted (right eye) retinas, respectively. Top row: Averaged peri-stimulus time histogram of retinal ganglion cell (RGC) spikes in response to 1-s full-field light stimulation at 12.84 log photons/cm²/sec, indicated by yellow bars from 10 to 11 s. Bottom row: The number of RGCs for each response type is shown in correlation with the channel position. (E–I) Representative immunostaining images 140–150 days after transplantation of NL-spheroids (top, E and F–I) and WL-spheroids (bottom, E). (E) RCVRN-positive (white) photoreceptors, many of which formed rosette structures, and SCGN-positive (red) cone bipolar cells were observed at the transplantation site. (F) Ribbon synapse formation between the dendritic tips of extending SCGN-positive (white) host cone bipolar cells and axon terminals of graft photoreceptors that form a rosette structure. CTBP2 (red) is expressed at the host-graft interface. (G) Graft photoreceptors expressing RHO (red) or OPN1LW/OPN1MW (white). (H and I) CALB1-positive (red, H) horizontal cells, CALB2-positive (white, H) amacrine or ganglion cells, and PRKCA-positive (red, I) bipolar rod cells were observed at the transplantation site. Scale bars: 500 μm (A), 1 mm (B), 100 μm (E, G–I), and 50 μm (F). NL-spheroid, non-layered spheroid; WL-spheroid, well-layered spheroid.

Immunohistochemical analysis was performed to confirm graft maturation and integration into the host retina 140–150 days after transplantation (graft DD 200–210). Transplanted NL- and WL-spheroids developed RCVRN-positive photoreceptors and SCGN-positive cone bipolar cells (top and bottom, respectively; Figure 5*E*). Some host cone bipolar cells extended dendrites toward graft photoreceptor rosettes, potentially forming synaptic connections, as indicated by presynaptic ribeye proteins (CTBP2) (Figure 5*F*). Most graft photoreceptors matured and formed rosette structures, as indicated by the expression of RHO (RetP1) and OPN1LW/OPN1MW (L/M-cone opsins) (Figure 5*G*). CALB1-positive HCs, CALB2-positive AC/RGCs, and PRKCA-positive rod bipolar cells were observed in the graft as previously reported ^3,10^ (Figures 5*H* and 5*I*).

## DISCUSSION

The present study identified a transient activation of WNT2B in a small sub-population, specifically in WL- spheroids, before the organization of the retinal layer structure, which is consistent with previous reports implicating canonical WNT ligands, including WNT2B, in vertebrate retinal development. ^6,25,26^ This small population expressing WNT2B was no longer detectable by the time when the phenotypic features of the NL and WL-spheroids became apparent at PS-D09 and beyond. Furthermore, we found that D6- Venus++/SSEA1- RPCs failed to self-organize into retinal layer structures at the initiation of spheroid formation in the absence of CHIR99021, a strong activator of canonical WNT signaling (Figure 1*J*). ^23^ These findings suggest that (1) canonical WNT signaling is required for retinal lamination, and (2) D3- Venus+++ RPCs may lack a responsive cell population with some WNT receptors for this process. Both scRNA-seq and immunohistological analyses demonstrated that cell populations expressing FZD7 receptors were absent in the D3-Venus+++ RPC populations (Figures 3*H* and 3*I*), implicating WNT2B– FZD7 signaling in the regulation of retinal layer formation. Because other canonical WNT ligands were not detected in PS-D05 WL-spheroids, WNT2B is a strong candidate for the initiation of layer formation, although we did not observe layer formation by the addition of WNT2B alone in our experimental setting.

Following the transient activation of WNT2B, distinct small cell populations expressing the ncWNT ligand WNT5A were observed in WL-spheroids. The ncWNT pathway comprises the planar cell polarity and calcium signaling pathways, which regulate cell polarity, promote cell migration and invasion, maintain stem cells, and modulate canonical WNT/β-catenin signaling. ^27,28^ Given these diverse and dynamic roles, WNT5A was initially considered a candidate for directly inducing the organization of retinal layer structures. However, the administration of WNT5A to D3-Venus+++ RPC-derived NL-spheroids from PS- D06 to PS-D12 failed to induce their differentiation into WL-spheroids, suggesting that some preceding process may be required to trigger layer formation before WNT5A. Nevertheless, transiently existing WNT5A-secreting RPC sub-populations in WL-spheroids may still be associated with this process, as suggested by the uniformly localized expression of WNT5A on the surface of WL-spheroids (Figure 4*H*). This implies that WNT5A may either play a supportive role in the formation of retinal layer structures or may be secondarily upregulated as a consequence of successful lamination.

While the final cell populations of WL- and NL-spheroids showed no substantial differences in scRNA-seq data, P-spheroids derived from D6-Venus+/++ cells exhibited three distinct cell populations: a RAX-positive CMZ-like population and two RAX-negative populations resembling RPE and spinal cord tissue, respectively. These three populations originated from D6-Rax::Venus+ cells, and the spinal cord-like tissue differentiated from RAX-positive cells, indicating unexpected cellular heterogeneity within this Rax::Venus+ population. These cell populations correspond to previously reported non-retinal off-target tissues of RO-derived grafts for transplantation, ^29^ supporting the notion that spheroids follow a differentiation process similar to that of ROs. Taken together, our results suggest that RAX-positive cells constitute a highly heterogeneous population of retinal progenitor and non-retinal lineage cells.

Unexpectedly, the present study revealed that the in vitro retinal layer structures on DD60 of retinal spheroids did not markedly influence structural or functional engraftment after transplantation into rats with retinal degeneration. Although well-aligned graft photoreceptor layers and their synaptic connections with host bipolar cells are generally considered essential for visual restoration, our results demonstrated that both types of spheroids—those with and without retinal layer structures in vitro but still consisting of very similar cell populations at the time of transplantation—enabled the maturation and layer formation of photoreceptors mostly in a rosette from within the graft. Additionally, these spheroid-derived photoreceptors made synaptic contact with host bipolar cells and elicited light responses in the host RGCs in MEA recordings. These results suggest that the host retinal environment strongly drives the alignment and integration of photoreceptors, even in NL-spheroids. Since NL- and WL-spheroids on PS- D35 shared very similar cell profiles in the scRNA-seq data, these given cell populations may result in similar functional and structural integration, regardless of the graft layer formulation in vitro.

In conclusion, this study provides new insights into the cellular and molecular mechanisms underlying the self-organization of retinal layer structures in hESC-derived RPCs. We identified the transient activation of WNT signaling pathways during the early phase of spheroid development, initially through canonical WNT2B–FZD7 signaling, followed by non-canonical WNT5A expression, which appears to sequentially contribute to the formation of retinal layer structures in vitro. However, our findings further demonstrate that even non-layer-forming RPC cells initially lacking WNT2B–FZD7 signaling at around DD40 can still achieve robust structural and functional integration following transplantation. Although intrinsic signaling and cellular heterogeneity play essential roles in early retinal morphogenesis in vitro, environmental factors within the host retina may ultimately guide the structural alignment and functional integration of the graft. This concept provides meaningful guidance for refining graft preparation strategies for efficient integration into future regenerative cell therapies.

## RESOURCE AVAILABILITY

### Lead contact

Further information and requests for resources and reagents should be directed to and fulfilled by the corresponding authors Michiko Mandai and Tomohiro Masuda.

### Materials availability

This study did not generate new unique reagents.

## Data and code availability

scRNA-seq data will be made available upon publication.

## Supporting information

Supplemental Informations

Supplemental CSV

## ACKNOWLEDGMENTS

We thank M. Matsumura and K. Kawai for their help with cell culture, Y. Ohigashi for immunohistochemistry, T. Yamada for MEA recordings, S. Yamasaki for his advice on cell culture and experiments, and J. Sho and T. Senba for supporting the animal experiments. We thank Editage (www.editage.jp) for the English language editing. The human ESC-KhES-1 cell line was provided by Kyoto University. This research was supported by the DECODE project in RIKEN, JSPS KAKENHI Grant Number JP21K09757, grants from the Manpei Suzuki Diabetes Foundation, and AMED grants JP13bm0204002, JP21bm0404073, and JP20bm0404024. This work was also supported by the Japan Science and Technology Agency, CREST (Grant Numbers JPMJCR21N6 and JPMJCR1926), the Medical Research Center Initiative for High Depth Omics, Multilayered Stress Diseases at TMDU, and TRIP-AGIS at RIKEN.

## AUTHOR CONTRIBUTIONS

Conceptualization: Y.I., T.M., and M.M.; Methodology: Y.I., T.M., I.N., and M.M.; Software: Y.I., M.Y., M.W., I.N., and M.M.; Formal Analysis: Y.I., M.Y., and M.W.; Investigation: Y.I., T.M., M.Y., M.W., I.N., and M.M.; Resources: Y.I., T.M., M.Y., M.W., K.N., M.F., M.T., and M.M.; Data Curation: Y.I., M.Y., M.W. and M.M.; Writing-Original Draft: Y.I.; Writing-Review & Editing: Y.I., T.M., M.Y., M.W., M.F., I.N., and M.M.; Visualization: Y.I., M.Y., and M.W.; Supervision: K.N., M.F., M.T., and M.M.; Project Administration: K.N., M.T., and M.M.; Funding Acquisition: M.T.; All authors reviewed and approved the final manuscript.

## SUPPLEMENTAL INFORMATION

**Document S1.** List of upregulated differentially expressed genes in clusters 22 and 23 on PS-D35 (related to Figure 2*G*). These genes were identified using the FindMarkers function implemented in Seurat. (Wilcoxon rank-sum test).

## EXPERIMENTAL PROCEDURES

### Human ESC maintenance

The Rax::Venus cell line (hESC-KhES-1) was provided by Kyoto University and used according to the hESC research guidelines of the Japanese government. This was maintained at RIKEN and Kobe City Eye Hospital as previously described. ^30^ Before passaging the cells, 6-well culture plates (Iwaki) were coated with iMatrix-511 (Nippi) in phosphate-buffered saline (PBS, Fujifilm Wako Pure Chemical) at 37 °C for 1 h. Human ESC colonies were treated with TrypLE Select Enzyme (Gibco) and 0.5 mM EDTA (Nacalai) at 37 °C for 8–10 min and dissociated by gentle pipetting. The dissociated cells were suspended in StemFit medium (AK03N, Ajinomoto) with 10 μM Y-27632 (Wako Pure Chemical Industries) and plated at a density of 1.0–1.3 × 10^4^ cells per well. The medium was replaced with StemFit medium without Y-27632 approximately 24 h after passage and replaced every day until the next passage.

### Retinal differentiation of hESCs

To prepare ROs, a modified SFEBq culture method with an induction-reversal culture method was used as described previously. ^15,17,18^ On the first day of SFEBq culture to DD0, recombinant human BMP4 (R&D Systems) was added to the culture at 1.5 nM on DD3 or DD6 (D3- and D6-BMP4), and its concentration was diluted by half of the medium every three days. Rax::Venus cell aggregates were cultured without BMP4 as a negative control for Venus-intensity (no-BMP4).

### Tissue dissociation for cell sorting and scRNA-seq

ROs on DD25 and spheroids on PS-D05, 09, 15, 21, and 35 were washed with PBS at room temperature (RT) and dissociated using dispersed solutions for neuronal cells (Fujifilm Wako Pure Chemical Corporation) at 37 °C for 30–70 min with gentle pipetting. After centrifugation and supernatant removal, the cell pellets were resuspended in Stain Buffer (PBS containing 2.0% fetal bovine serum) and filtered through a 40-μm cell strainer (pluriSelect) to remove cell clumps. Cells were counted for scRNA-seq using a hemocytometer, and cell viability was calculated. Dissociated cells were preserved in Stain Buffer at 4 °C in the dark until the next step.

### FACS procedures

FACS was performed using a FACS Aria II flow cytometer (BD Biosciences). FACSDiva or FlowJo software (BD Biosciences) was used for the analysis (Figures 1*A–D*). Dead cells, cell debris, and cell multiplets were excluded from the analysis and sorted by gating with forward and side scatters as indicators. The cells were sorted into CD15-BV421-negative fractions with Rax::Venus+++, Rax::Venus++, or Rax::Venus+/++ fractions. The cells were collected in tubes containing neural retinal differentiation medium for the spheroid assay. After FACS, sorted cells were examined for fluorescence intensity (Figure 1*D*).

### Retinal spheroid formation and differentiation

After cell sorting, single cells in Stain Buffer were centrifuged, and the supernatants were removed. Cell pellets were resuspended with NR differentiation medium supplemented with 10 μM Y-27632 (Wako Pure Chemical Industries), 3 μM CHIR99021 (Bio Vision), 300 nM SAG (Enzo Biochem), and 100 ng/mL human recombinant FGF8 (Fujifilm Wako Pure Chemical) and counted with Countess II (WO/2023/090427). The cells were plated in a 96-well plate at a density of 12000 cells/well and cultured at 37 °C in a humidified atmosphere containing 5% CO_2_. The medium was replaced every three days. As for administration tests, human recombinant WNT2B (Abnova) was added to the medium at 100 and 500 ng/μll from PS-D0 to PS-D6, and human recombinant protein WNT5A (R&D Systems) was added to the medium at 50, 200, 500, and 1000 ng/μll from PS-D12 to PS-D18.

### Single-cell RNA-seq analysis

Single-cell RNA sequence libraries were prepared using the 10X Genomics Chromium 3’ v3.1 library kit and then sequenced on an Illumina HiSeq X. FASTQ files were aligned to GRCh38 v. 2020-A using 10X Genomics CellRanger v6.1.1. Quality control metrics, such as UMI count, gene count, and mitochondrial percentage, were assessed, and cells outside a defined range of feature counts and mitochondrial percentages were excluded. A mean of 10,376 cells, ranging from 7,020 to 12,418 cells, were included in each original sample, and a mean of 10,268 cells, ranging from 6,960 to 12,261 cells, were included after filtering. Seurat v4.3.0 was used for normalization, feature selection (vst, number of features = 2000), and downstream analyses based on the previously reported workflow tutorial. ^31^ In the downstream analysis, 1) filtered data of WL-, NL-, and P-spheroids on PS-D35 were merged into one Seurat object, and 2) filtered data of WL- and NL-spheroids on PS-D05, -D09, and -D15 were merged into one at each time point. The data were then scaled, principal components were computed, nearest neighbors were determined, and UMAP embedding was performed. ^31^ Cell-cycle bias correction was performed using the CellCycleScoring function implemented in Seurat. Cell clusters were identified and manually merged based on previously identified markers of RPCs, RGCs, photoreceptors, ACs/HCs, and RPE cells. ^19,20^ DEGs from the merged PS-D35 data were calculated using the FindMarkers function implemented in Seurat and used for gene ontology analysis using Metascape v3.5. To analyze the merged data of WL- and NL-spheroids on PS-D05, -D09, and -D15, CellChat v1.5.0 was used to analyze ligand-receptor interactions between individual cells. ^22^

### Transplantation into nude rats

All animal experiments were performed according to the ARVO statement for the use of animals in ophthalmic and vision research and approved by the Animal Research Committee at the RIKEN Center for Biosystems Dynamics Research. SD-Foxn1 Tg(S334ter) 3LavRrrc nude rats, which are an immunodeficient model of progressive retinal degeneration, were obtained from the Rat Resource and Research Center. ^24^ For transplantation, rats were anesthetized with medetomidine hydrochloride (0.375 mg/kg), midazolam (2 mg/kg), or butorphanol tartrate (at least 2.5 mg/kg). Retinal sheets were manually cut from WL- and NL-spheroids on DD60 and inserted into the subretinal space of rats aged approximately 6 months, as previously described. ^3,10,17^ Transplanted rats were sacrificed, and retinas were harvested via isoflurane inhalation after 10–11 months for MEA and immunohistochemistry.

### Electrophysiology: MEA recordings

The following procedure was performed under a dim LED light with a peak wavelength of 690 nm. After 24 h of dark adaptation, animals were anesthetized with isoflurane and euthanized by cervical dislocation. The eyes were enucleated, and the cornea and vitreous were removed to isolate the retina. The transplanted regions were identified visually, and the retina was trimmed around the graft area. The tissue was placed with the ganglion cell (RGC) side attached to a 60-electrode MEA (Multi Channel Systems, 60pMEA200/30iR-Ti; electrode size: 30 × 30 μm; spacing: 200 μm) by using mild suction and a mesh-covered anchor placed on top of the retina. The retina was continuously perfused with warmed (34 °C) oxygenated (95% O_2_/ 5% CO_2_) Ames’ medium (Sigma, A1420) at 3–3.5 mL/min. The recordings began 40 min after the start of the perfusion. The extracellular voltage signals were amplified and digitized at 20 kHz using a USB-ME64-System (Multi Channel Systems).

Full-field light stimuli (12.84 log photons/cm²/s, corresponding to approximately 6.5 lux or 3.4 × 10D Rh∗/rod/s) of 1-second duration were delivered under the dark background condition using a white LED (NSPW500C, Nichia Corp., Tokushima, Japan). The LED output was modulated using an electronic pulse generator (DSP-420, Dia-Medical System, Tokyo, Japan) controlled by the TTL signals from the MEA system.

### Immunohistological procedures

Spheroids on DD60 and freshly harvested rat eyes after transplantation were fixed with 4% paraformaldehyde (Fujifilm Wako Pure Chemical Corporation), infiltrated with 30% sucrose/PBS overnight at 4 °C, embedded in O.C.T. compound (Sakura Finetek Japan), and stored at −30 °C. Cryo-sections of 10 μm thickness for spheroids and 50 μm thickness for the transplanted rat eyes were performed.

Samples were washed with PBS and blocked with a blocking solution of 1% bovine serum albumin and 0.1% Tween 20 in PBS for 1 h at RT for 10 μm-thick cryo-sections and overnight at 4 °C for 50 μm-thick cryo-sections. Primary antibodies were diluted appropriately (Table S1) with a blocking solution and incubated with 10-μm-thick samples overnight and 50-μm-thick samples for three days at 4 °C. Samples were then washed three times with 0.05% Tween-20 in PBS, and the appropriate secondary antibodies and DAPI (Molecular Probes) were incubated with 10-μm-thick samples for 1 h at RT and 50-μm-thick samples for three days at 4 °C. After secondary antibody staining, the samples were washed thrice with 0.05% Tween-20 in PBS and mounted using FluorSave Reagent (Merck Millipore). Labeled samples were kept in the dark at 4 °C before imaging with a BZ-X800 Keyence Fluorescence Microscope or a confocal microscope (Zeiss LSM700 and 980 and Leica-TCS SP8).

