## Supplemental Informations for "Genomic characterization of sub-populations in human pluripotent stem cell-derived retinal progenitor cells that drive retinal layer structure"

**Supplemental Table**

**Table S1.** List of antibodies used for immunohistochemical analysis, related to Experimental procedures.

| Target | Antibody Name | Host | Dilution | Vendor | Catalog Number |
|---|---|---|---|---|---|
| BRN3 | anti-Brn-3 | Goat | 1:1000 | Santa Cruz | sc-6026 |
| CALB1 | anti-Calbindin D-28K | Rabbit | 1:500 | Sigma-Aldrich | ABN2192 |
| CALB2 | anti-Calretinin, clone 6B8.2 | Mouse | 1:1000 | Sigma-Aldrich | MAB1568 |
| CTBP2 | anti-CtBP2 | Mouse | 1:500 | BD Biosciences | 612044 |
| CRX | anti-CRX (clone 4G11) | Mouse | 1:20000 | Abnova | H00001406-M02 |
| EMX2 | anti-EMX2 | Rabbit | 1:500 | invitrogen | PA5-34415 |
| FZD7 | anti-Frizzled 7 | Rabbit | 1:500 | Abcam | Ab64636 |
| Human Nuclei | anti-Nuclei (clone 235-1) | Mouse | 1:1000 | Sigma-Aldrich | MAB1281 |
| OPN1LW/OPN1MW | anti-opsin, red/green | Rabbit | 1:1000 | Sigma-Aldrich | AB5405 |
| PAX6 | anti-human Pax-6 | Mouse | 1:500 | BD Biosciences | 561462 |
| PRKCA | anti-protein kinase Cα | Goat | 1:500 | R&D systems | AF5340 |
| RCVRN | anti-recoverin | Rabbit | 1:1000 | Proteintech | 10073-1-AP |
| RHO | anti-opsin (clone RET-P1) | Mouse | 1:1000 | Sigma-Aldrich | O4886 |
| SCGN | anti-secretagogin | Sheep | 1:2000 | Bio Vendor | RD184120100 |
| VSX2 | anti-Chx10 (visual system homeobox 2) | Sheep | 1:1000 | Exalpha | X1180P |
| WNT5A | anti-Wnt-5a (A-5) | Mouse | 1:100 | Santa Cruz | sc-365370 |
| ZIC1 | anti-human/mouse ZIC1 | Goat | 1:200 | R&D systems | AF4978 |

### Supplemental Figures

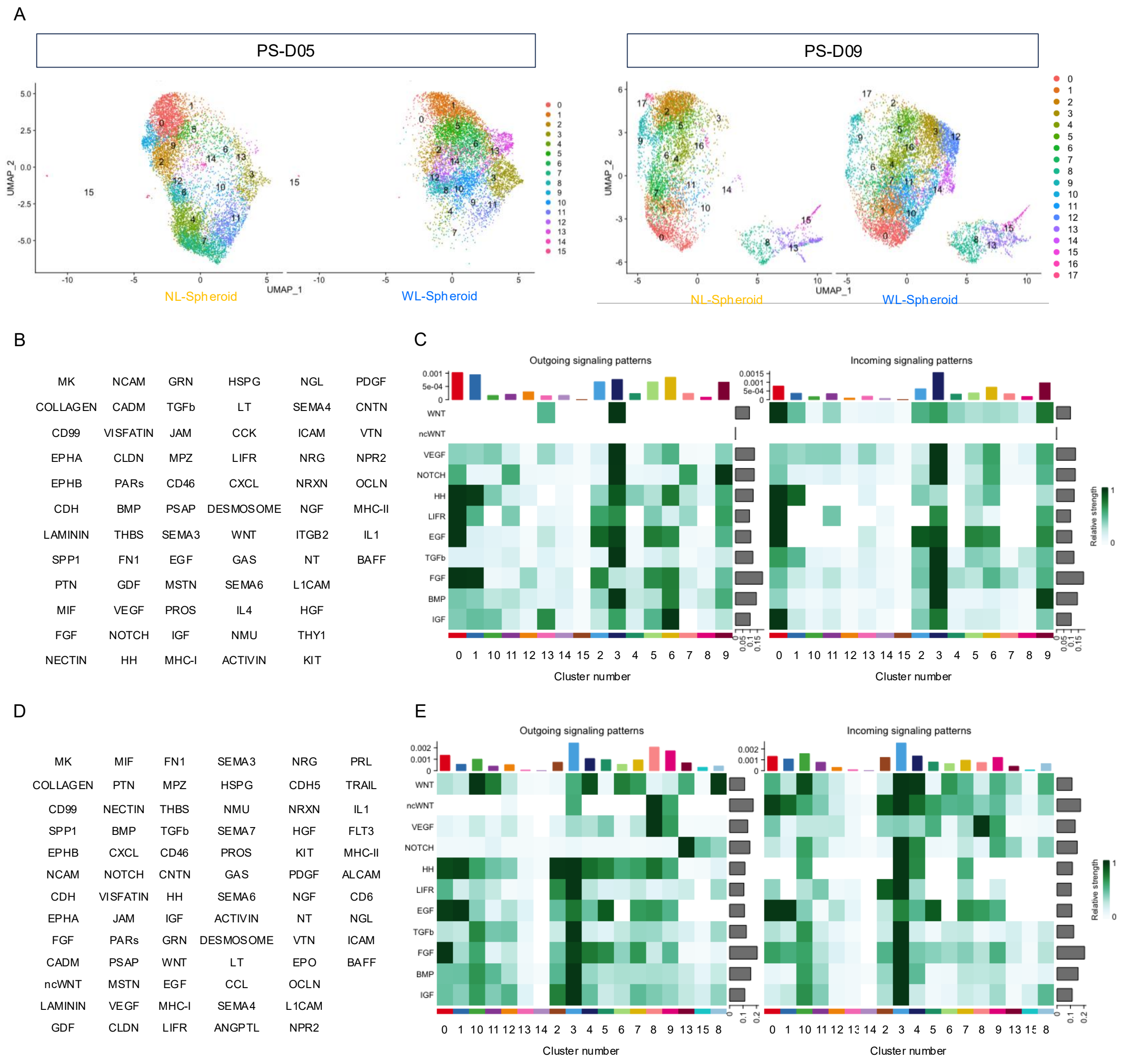


**Figure S1. Significantly activated signaling pathways on PS-D05 and -D09**

(A) UMAPs of merged single cell data of NL- and WL-spheroids on PS-D05 and PS-D09 (left and right, respectively)
(B) Potentially significant pathways on PS-D05, calculated using CellChat. Sixty-eight pathways were detected as significantly activated signaling pathways.
(C) Heatmaps of inferred signaling networks of typical pathways in retinal development on PS-D05. Outgoing and incoming signaling means ligand and receptor sides, respectively.
(D) Potentially significant pathways on PS-D09, calculated using CellChat. Seventy-five pathways were detected as significantly activated signaling pathways.
(E) Heatmaps of inferred signaling networks of typical pathways in retinal development on PS-D09. Outgoing and incoming signaling means ligand and receptor sides, respectively.

A

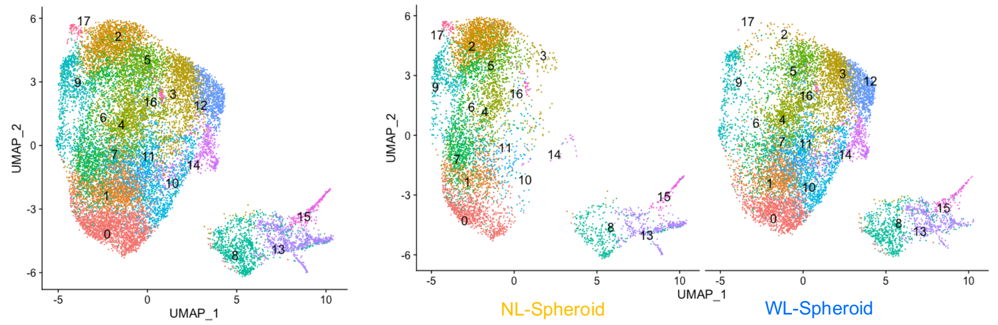


| | 0 | 1 | 2 | 3 | 4 | 5 | 6 | 7 | 8 | 9 | 10 | 11 | 12 | 13 | 14 | 15 | 16 | 17 |
|---|---|---|---|---|---|---|---|---|---|---|---|---|---|---|---|---|---|---|
| NL | 850 | 597 | 1403 | 116 | 477 | 469 | 950 | 435 | 339 | 486 | 35 | 205 | 0 | 340 | 24 | 126 | 40 | 68 |
| WL | 1211 | 979 | 93 | 1372 | 955 | 723 | 174 | 588 | 631 | 426 | 859 | 686 | 834 | 405 | 487 | 123 | 40 | 1 |

B

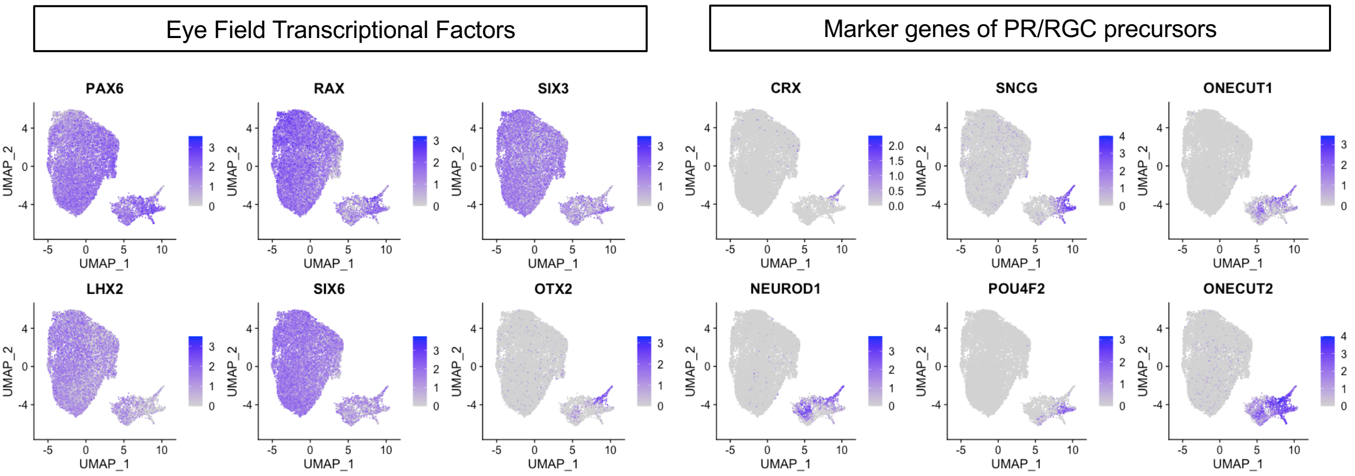


**Figure S2. Merged single cell RNA-seq data of NL- and WL-spheroids on PS-D09**

(A) UMAP of merged single cell data of NL- and WL-spheroids on PS-D09. Spheroids consisted of two groups of clusters.
(B) Feature plots showing the expression of eye field transcriptional factors and marker genes of photoreceptor and retinal ganglion precursor cells in blue.
